# tinyRNA: precision analysis of small RNA-seq data with user-defined hierarchical selection rules

**DOI:** 10.1101/2022.06.29.497994

**Authors:** Alex J. Tate, Kristen C. Brown, Taiowa A. Montgomery

## Abstract

**Summary:** tinyRNA performs precision analysis of small RNAs, including miRNAs, piRNAs, and siRNAs, from high-throughput sequencing experiments. At the core of tinyRNA is a highly flexible counting utility, tiny-count, that allows for hierarchical assignment of small RNA reads to features based on positional information, extent of feature overlap, 5’ nucleotide, length, and strandedness. tinyRNA provides an all-in-one solution for small RNA-seq data analysis, with documentation and statistics generated at each step for accurate, reproducible results.

**Availability and Implementation:** tinyRNA tools are implemented in Python and R, and the pipeline workflow is coordinated with CWL. tinyRNA is free and open-source software distributed under the GPLv3 license. tinyRNA is available at https://github.com/MontgomeryLab/tinyRNA.

**Supplementary information:** Reference data, including genome sequences and features tables, for certain species can be found at https://www.MontgomeryLab.org.

## INTRODUCTION

Small RNAs interact with larger RNAs, such as mRNAs, through complementary base-pairing to regulate gene expression. There is tremendous diversity amongst small RNAs both within and between species, but they are unified by their association with Argonaute/Piwi proteins (Ghildiyal and Zamore, 2009). Genetic requirements, particularly Argonautes, are typically used to classify small RNAs, although other features, such as mode of action and genomic origin, can also distinguish or further subclassify small RNAs. High-throughput sequencing is a widely used tool for discovery and analysis of small RNAs. However, the diversity and complexity of small RNA pathways presents several computational challenges when classifying and analyzing the reads generated by high-throughput sequencing. For example, small RNA libraries are often contaminated with decay intermediates of longer RNAs, such as rRNAs and mRNAs. These contaminants can be difficult to distinguish from authentic small RNAs produced from the same features. Additionally, small RNAs are subjected to decay and regulatory modifications, such as the addition of 3’ untemplated nucleotides, that can functionally distinguish them from their unmodified counterparts.

Further complicating data analysis, distinct classes of small RNAs, such as piwi-interacting RNAs (piRNAs) and small interfering RNAs (siRNAs), can be produced from the same locus (Kawamura, et al., 2008) (Das, et al., 2008). The tinyRNA project address these challenges with tiny-count, a precision counting tool designed specifically for small RNAs, and provides an end-to-end workflow that is accurate, reproducible, portable, and utilizes best practices in small RNA high-throughput sequencing data analysis. The project is open-source and community driven.

## METHODS

### Overview

Installation of tinyRNA and its dependencies is automated with an installation script that places all components within an isolated conda environment. Prior to running tinyRNA, users specify their sample information and feature selection rules in csv-formatted template files. tinyRNA execution begins with the automated generation of a workflow in Common Workflow Language (CWL) (Crusoe, et al., 2021) configured for the input parameters provided by the user. Preprocessing of FASTQ files, including adapter trimming and quality filtering, is performed by fastp (Chen, et al., 2018). Unique sequences are counted and collapsed by tinyRNA’s collapser utility, tiny-collapse, substantially reducing the resource demands of genomic alignment and feature counting. tiny-collapse can also trim the degenerate bases often included in the adapter sequences used in library preparation. Collapsed reads are then aligned to a reference genome using bowtie (Langmead, et al., 2009) and a reference index or genome sequence provided by the user. Next, tiny-count evaluates these alignments for feature assignment while utilizing the Genomic Array of Sets and GFF reader from HTSeq (Anders, et al., 2015). Feature assignment is based on the user’s selection rules and the features defined in their GFF/GTF formatted file inputs. Any number of rules can be used to classify reads by sequence attributes such as 5’ nucleotide, length, and strandedness. The read count for each sequence is normalized by the numbers of genomic hits and feature matches before assignment to features. If the data contains biological replicates, differential expression analysis is then automatically performed by tiny-deseq, a wrapper around DESeq2 (Love, et al., 2014). Finally, the outputs of tiny-count and tiny-deseq are used to produce publication-ready plots with tinyRNA’s plotter utility, tiny-plot. The outputs of each run, including quality reports, processed data, mapping and assignment statistics, counts tables, size plots, class charts, and a variety of scatter plots, are placed in a timestamped directory with full run documentation (Figure 1).

**Figure 1.**
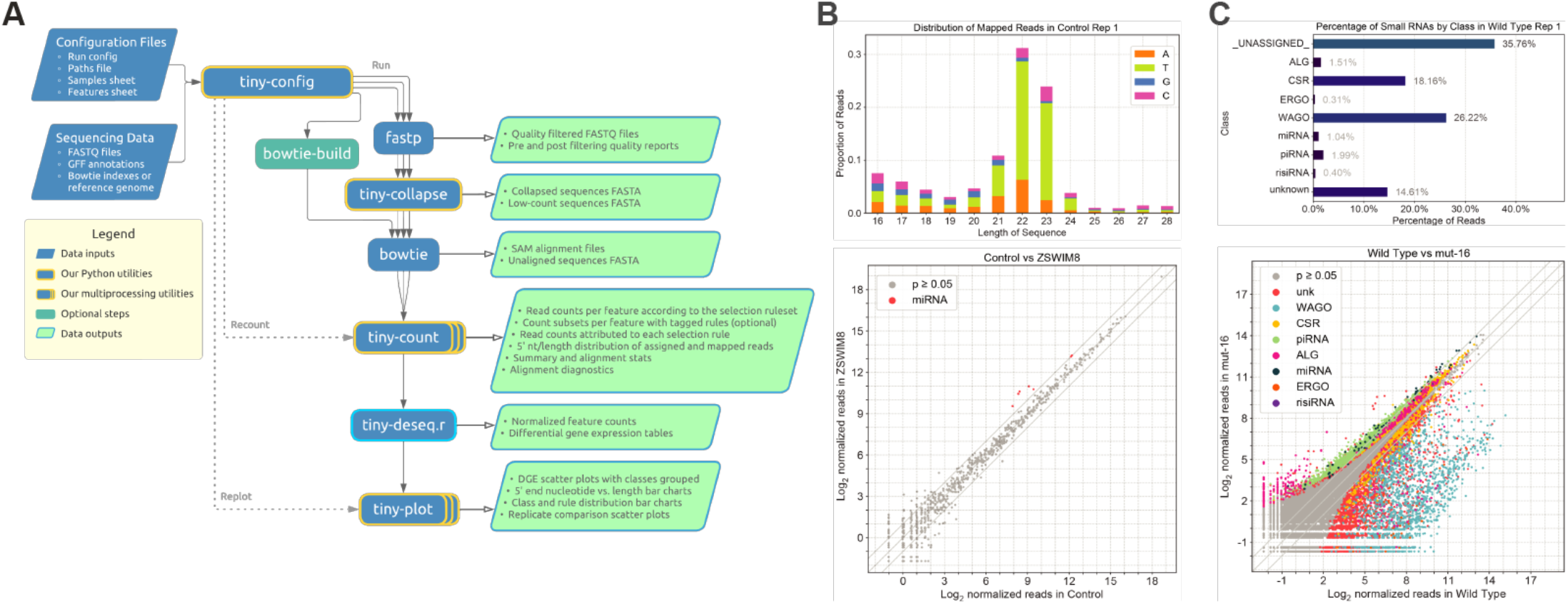
tinyRNA workflow. (A) tinyRNA flowchart. (B-C) A selection of graphical outputs from a human miRNA analysis (Shi, et al., 2020) (B), and a *C. elegans* total small RNA analysis (Reed, et al., 2020) (C).

### Counting Precision

The tiny-count utility provides precise control over the attribution of reads at alignment loci with user-defined selection rules. The selection process resolves ambiguities in feature assignment at loci with multiple overlapping features and the associated loss of feature count precision. It also allows for reads from distinct classes of small RNAs produced from the same feature or genomic interval to be treated separately at counting, plotting, and differential expression analysis steps. Feature selection and read filtering occurs in three stages. In stage 1, features of interest are retrieved from GFF/GTF-formatted input files based on their attributes such as small RNA classification or gene biotype. In stage 2, these features undergo hierarchical elimination followed by evaluation of their extent of overlap with each read alignment. Selection rules can include partial, full, or exact overlap. Additionally, features can be filtered based on whether their 5’ or 3’ ends are anchored to the alignment. Remaining candidate features move to stage 3 where they are filtered based on the alignment’s sequence attributes, such as length, strandedness relative to the feature, and 5’ nucleotide. A hierarchical selection scheme allows users to assign reads preferentially or uniformly to one set of features or another. For example, reads aligning to rRNAs can be excluded from any or all other features by assigning rRNAs to a lower hierarchy value (closer to or equal to 1) than other features. Features which pass the selection process are assigned a normalized portion of the counts associated with the alignment sequence. For a sequence with *n* read counts and *m* alignments, a maximum of *n*/*m* reads can be assigned to features at each locus. If *k* features pass selection at one of these loci, each of these features will be assigned 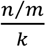 reads.

### Performance

Most steps in tinyRNA run concurrently across libraries to minimize runtime. A user-defined global thread count is set for fastp and bowtie to utilize multi-core computing resources. Performance-critical sections of tinyRNA components have been carefully written to optimize execution at the bytecode level. Runtime will vary considerably depending on the number of genomic alignments produced and the characteristics of input sample files and parameters. However, we have found that tiny-count typically processes alignment files at a rate of 3 × 10^5^ alignments per second. Building a bowtie index can add significant time to the analysis, particularly for large genomes. For this reason, tinyRNA automatically amends configuration files so that bowtie indexes are only built during the first analysis for a particular genome.

### Convenience and Reproducibility

tinyRNA is designed for users whose backgrounds span computational naivety to programming comprehension. Configuration files are modular and reusable rather than residing in a single monolithic file. This allows the entire collection of utilities to be executed with a single command whose sole argument is the primary configuration file. A standard small RNA analysis configuration that follows best practices in the field is provided as a modifiable template in YAML format. Copies of configuration files are placed in timestamped output directories that are organized into subdirectories by step to simplify documentation and collaboration. Most tinyRNA components are written in Python using object-oriented design patterns for transparency and easy modification by advanced users. tinyRNA can also be extended to include any number of intermediate steps by modifying the workflow CWL. These advanced modifications are permissible by the installation script for easy integration.

## SUMMARY

tinyRNA automates precision analysis of small RNA high-throughput sequencing data following best practices and produces complete run documentation for accuracy and reproducibility. It is compatible with data from any species containing a reference genome or transcriptome. tinyRNA can accommodate relatively simple data analyses, such as miRNA expression in human cell lines, to complex analyses of multiple classes of small RNAs from whole organisms, such as *C. elegans*.

## ACKNOWLEDGEMENTS

This work was supported by the National Institutes of Health [R35GM119775 to T.A.M.].

## REFERENCES

Anders, S., Pyl, P.T. and Huber, W. HTSeq--a Python framework to work with high-throughput sequencing data. Bioinformatics 2015;31(2):166–169.

Chen, S., et al. fastp: an ultra-fast all-in-one FASTQ preprocessor. Bioinformatics 2018;34(17):i884–i890.

Crusoe, M.R., et al. Methods Included: Standardizing Computational Reuse and Portability with the Common Workflow Language. Communications of the ACM 2022;65(6):54–63.

Das, P.P., et al. Piwi and piRNAs act upstream of an endogenous siRNA pathway to suppress Tc3 transposon mobility in the Caenorhabditis elegans germline. Mol Cell 2008;31(1):79–90.

Ghildiyal, M. and Zamore, P.D. Small silencing RNAs: an expanding universe. Nat Rev Genet 2009;10(2):94–108.

Kawamura, Y., et al. Drosophila endogenous small RNAs bind to Argonaute 2 in somatic cells. Nature 2008;453(7196):793–797.

Langmead, B., et al. Ultrafast and memory-efficient alignment of short DNA sequences to the human genome. Genome Biol 2009;10(3):R25.

Love, M.I., Huber, W. and Anders, S. Moderated estimation of fold change and dispersion for RNA-seq data with DESeq2. Genome Biol 2014;15(12):550.

Reed, K.J., et al. Widespread roles for piRNAs and WAGO-class siRNAs in shaping the germline transcriptome of Caenorhabditis elegans. Nucleic Acids Res 2020;48(4):1811–1827.

Shi, C.Y., et al. The ZSWIM8 ubiquitin ligase mediates target-directed microRNA degradation. Science 2020;370(6523).

